# Exercise twice-a-day potentiates markers of mitochondrial biogenesis in men

**DOI:** 10.1101/547489

**Authors:** Victor A. Andrade-Souza, Thaysa Ghiarone, Andre Sansonio, Kleiton Augusto Santos Silva, Fabiano Tomazini, Lucyana Arcoverde, Jackson Fyfe, Enrico Perri, Nicholas Saner, Jujiao Kuang, Romulo Bertuzzi, Carol Gois Leandro, David J. Bishop, Adriano E. Lima-Silva

**Affiliations:** Department of Physical Education and Sports Science, Academic Center of Vitoria, Federal University of Pernambuco, Vitoria de Santo Antao, Pernambuco, Brazil; Department of Medicine, University of Missouri School of Medicine, Columbia, Missouri, USA; School of Exercise and Nutrition Sciences, Faculty of Health, Deakin University, Burwood, Victoria, Australia; Institute for Health and Sport, Victoria University, Melbourne, Victoria, Australia; Endurance Performance Research Group, School of Physical Education and Sport, University of São Paulo, Sao Paulo, Brazil; School of Medical and Health Sciences, Edith Cowan University, Joondalup, Western Australia, Australia; Human Performance Research Group, Academic Department of Physical Education, Technological Federal University of Parana, Curitiba, Parana, Brazil

**Keywords:** muscle glycogen, high-intensity exercise, mitochondrial biogenesis, molecular signalling, transcription factor

## Abstract

Endurance exercise begun with reduced muscle glycogen stores seems to potentiate skeletal muscle protein abundance and gene expression. However, it is unknown whether this greater signalling responses is due to performing two exercise sessions in close proximity - as a first exercise session is necessary to reduce the muscle glycogen stores. In the present study, we manipulated the recovery duration between a first muscle glycogen-depleting exercise and a second exercise session, such that the second exercise session started with reduced muscle glycogen in both approaches but was performed either 2 or 15 h after the first exercise session (so-called “twice-a-day” and “once-daily” approaches, respectively). We found that exercise twice-a-day increased the nuclear abundance of transcription factor EB (TFEB) and nuclear factor of activated T cells (NFAT) and potentiated the transcription of peroxisome proliferator-activated receptor-□ coactivator 1 alpha (PGC-1α), peroxisome proliferator-activated receptor alpha (PPARα) and peroxisome proliferator-activated receptor beta/delta (PPARβ/δ) genes, in comparison with the once-daily exercise. These results suggest that part of the elevated molecular signalling reported with previous “train-low” approaches might be attributed to performing two exercise sessions in close proximity. The twice-a-day approach might be an effective strategy to induce adaptations related to mitochondrial biogenesis and fat oxidation.

## INTRODUCTION

Endurance exercise is a powerful stimulus affecting cytoplasmic and nuclear proteins, and genes encoding mitochondrial proteins, with a subsequent increase in mitochondrial biogenesis (i.e., the generation of new mitochondrial components leading to increased mitochondrial content and respiratory function) (1–8). While these responses are affected by the nature of the exercise (e.g., the exercise intensity (2, 9)), there is evidence substrate availability is also a potent modulator of this response (10–13). It has been hypothesized that initiating endurance exercise with low muscle glycogen stores (the so-called “train-low” approach) results in a greater increase in the content of nuclear proteins (14, 15) and in the transcription of genes (16–19) associated with mitochondrial biogenesis. However, although the “train-low” strategy has been reported to potentiate skeletal muscle signalling responses related to mitochondrial biogenesis (16, 20–22), there are also contrasting findings showing no effects (23, 24) and a consensus is yet to be reached.

Some of the evidence supporting the “train-low” approach is based on performing a first exercise session to reduce muscle glycogen stores, which is followed by a second exercise session 1 to 3 h later – the so-called “twice-a-day” approach (20–23, 25–27). Although the second exercise session will start with reduced muscle glycogen stores, it is difficult to determine if any changes are due to performing the second exercise session with low muscle glycogen or performing the second exercise session soon after the first. The transcriptional responses of many genes associated with mitochondrial biogenesis peak approximately 3 h post exercise and return to basal levels within 8 to 12 h (5, 13, 28–31), including the transcription of peroxisome proliferator-activated receptor-□ coactivator 1 alpha (PGC-1α) - considered a key regulator of mitochondrial biogenesis (32). Thus, it is possible that reported increases in gene expression with the twice-a-day approach can be attributed to performing the second exercise session close to the first, when there is an already increased expression of genes associated with mitochondrial biogenesis.

The aim of this study was to investigate whether greater exercise-induced signalling with the “train-low” approach can be attributed to the cumulative effects of performing two exercise sessions in close proximity. In the present study, we manipulated the recovery duration between a first muscle glycogen-depleting exercise and a second exercise session (i.e., a “once-daily” *vs*. a “twice-a-day” approaches). In the once-daily condition, muscle glycogen content was reduced via evening exercise (prolonged exercise) followed by a carbohydrate-restricted period, and a second exercise (i.e., high-intensity interval exercise) was performed the next day (i.e., 15 h between exercise sessions). In the twice-a-day condition, the same exercises were used but with a short recovery period between exercise sessions (i.e., 2 h between exercise sessions). A high-intensity interval exercise session undertaken without a prior muscle glycogen-depleting exercise served as a control. We hypothesized that the “twice-a-day” approach would induce a more elevated molecular signalling response during the second exercise session, compared to the “once-daily” approach.

## MATERIALS AND METHODS

### Participants

Eight healthy men [age: 30.8 ± 3.5 years, body mass: 78.7 ± 9.9 kg, height: 1.76 ± 0.07 m, body fat: 13.6 ± 5.1%, peak oxygen uptake 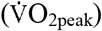: 37.1 ± 6.4 mL·kg^-1^·min^-1^, peak power derived from a graded exercise test: 229.6 ± 38.2 W, and first and second lactate thresholds: 78.1 ± 20.9 and 170.0 ± 37.7 W, respectively] participated in this study. Participants were recruited by word-of-mouth conversations, online postings, and flyers posted at the University Campus. Participants were considered eligible if were physically active (more than 3 times/week) and accustomed to cycling. Participants were excluded if they had history or signs of cardiac, metabolic, or respiratory diseases and/or had been using any prescription medications. A medical doctor administered a medical history screening tool for this purpose. Participants were informed about the procedures, risks, and benefits associated with the protocol, before they signed a consent form agreeing to participate in this study, which was approved by the Research Ethics Committee of Federal University of Pernambuco (approval number: 30378414.8.0000.5208). The study was conducted according to the principles presented in the Declaration of Helsinki.

Sample size was calculated using a G*Power software (Heinrich-Heine-University Düsseldorf, version 3.1.9.2, Düsseldorf, Germany), assuming an effect of exercising with reduced muscle glycogen content on PGC-1α mRNA expression of 1.34 (16), an alpha of 0.05, and a desired power of at least 0.80. We used PGC-1α mRNA expression as the main primary outcome as this gene has been considered a key regulator of mitochondrial biogenesis (32). The effective sample size necessary to achieve statistical significance was 6 participants; however, sample size was increased to 8 participants to account for potential dropouts or insufficient muscle sample size due to technical problems.

### Study overview

Each participant completed three experimental trials in a crossover, randomized, and incomplete balanced Latin Square counterbalanced measure design. The randomization was performed using software available at http://www.randomized.org and controlled by an investigator of our laboratory who was not conscious of the aim of the study. All tests were performed at Vitoria Santo Antao Campus of Federal University of Pernambuco from March to November 2015.

An overview of the experimental design is shown in Fig. 1. Briefly, in the once-daily approach participants performed the muscle glycogen-depleting exercise in the evening (2000 – 2200 h) followed by an overnight fast. The next morning (0800 h), participants ate a low-carbohydrate breakfast (carbohydrate: 42.7 ± 5.0 kcal, 7%; fat: 365.9 ± 43.1 kcal, 60%; protein: 201.3 ± 23.7 kcal, 33%) and then performed the high-intensity interval exercise session in the afternoon (1300 h). In the twice-a-day approach, participants ate a low-carbohydrate breakfast on the morning of the experimental day (0800 h), and then performed a muscle glycogen-depleting exercise (0900-1100 h), followed by a 2-h rest period and the high-intensity interval exercise (1300 h). In the control condition, participants consumed the same low-carbohydrate breakfast as the two other experimental trials (0800 h), and then performed the same high-intensity interval exercise session in the afternoon (i.e., 1300 h). Skeletal muscle biopsies from the *vastus lateralis* and venous blood samples were taken before, immediately after, and 3 h after completion of the high-intensity interval exercise sessions. Participants were not allowed to eat any food between the breakfast and the last muscle biopsy in all experimental trials. Water was provided ad libitum throughout each trial. Each experimental trial was separated by approximately two weeks to washout any residual effect of fatigue or damage caused by exercise and the multiple muscle biopsies. The main primary outcome measure was PGC-1α mRNA, while the second outcome measures were nuclear/cytosolic proteins and other genes associated with mitochondrial biogenesis.

**Figure 1.**
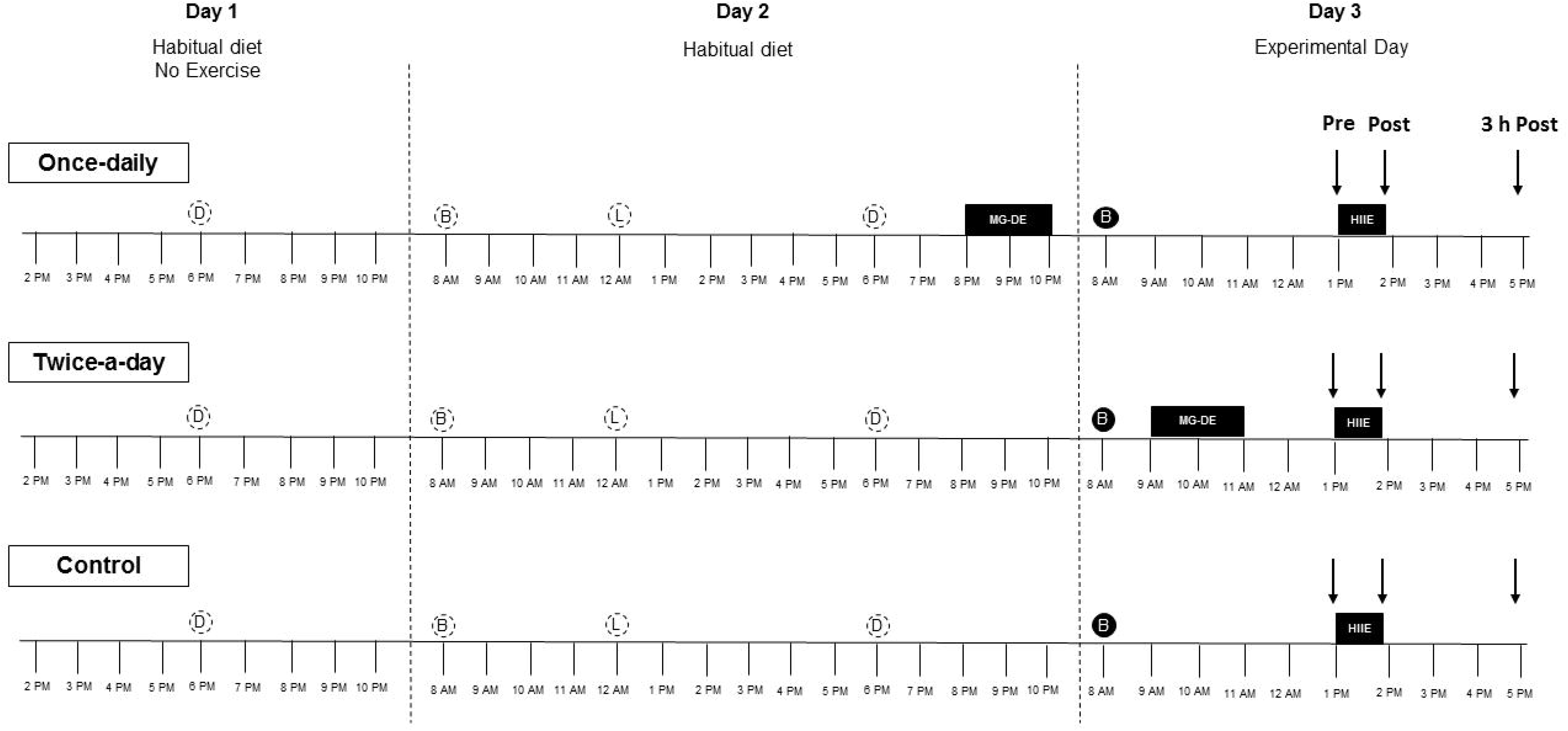
Experimental design. D, dinner; B, breakfast; L, lunch; MG-DE, muscle glycogen-depleting exercise; HIIE, high-intensity interval exercise. Open dashed circles indicate that participants replicated their usual diet, while closed circles indicate that participants ate a low-carbohydrate (CHO) breakfast [CHO: 42.7 ± 5.0 kcal (~7%), fat: 365.9 ± 43.1 kcal (~60%), protein: 201.3 ± 23.7 kcal (~33%)]. Black arrows indicate time point whenre muscle biopsies and blood samples were taken.

### Exercise protocols

#### Preliminary test

One week prior to the commencement of this study, participants performed a graded exercise test to volitional fatigue on a cycloergometer (Ergo-Fit 167, Pirmasens, Germany). The test commenced at 50 W, and thereafter intensity was increased by 25 W every 4 min, with a 1-min break between stages, until volitional exhaustion (33). The test was interrupted when the participant could no longer maintain the required cadence (70 rpm). Strong verbal encouragement was provided to each participant.

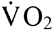 was measured breath-by-breath throughout the test using an automatic analyzer (Cortex, Metalyzer 3B^®^, Saxony, Germany). Before each test, the gas analyzer was calibrated using ambient air and a cylinder of known gas concentration (12% O_2_ and 5% CO_2_). The volume was calibrated using a 3-L syringe (Quinton Instruments, Washington, US). Capillary ear lobe blood samples were taken at rest and immediately after each 4-min stage of the test for determination of plasma lactate concentration. The first lactate threshold was visually identified by two experienced investigators as the first increase in plasma lactate concentration above resting level. The second lactate threshold was calculated by the modified Dmax method (34). This was determined by the point on the polynomial regression curve that yields the maximal perpendicular distance to the straight line connecting the first lactate threshold and the final stage of the test. 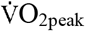 was defined as the highest 30-s average 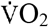 during the test and peak power was determined as the highest workload reached. If a participant did not complete the final 4-min stage, then the peak power was determined using the following equation:

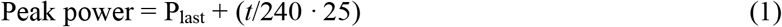

where P_last_ is the power in Watts of the last completed stage performed by the participant, *t* is the time (in seconds) sustained during the last incomplete stage, and 25 corresponds to the increments in power (Watts) at each stage.

#### Muscle glycogen-depleting exercise

To reduce muscle glycogen stores, participants cycled for 100 min at a power corresponding to 50% of the difference between their first and second lactate threshold (124 ± 27 W, 54 ± 5% of peak power). Then, after an 8-min rest, participants performed six 1-min exercise bouts at 125% of their peak power (287 ± 46 W) interspersed with 1-min rest periods (35). This protocol has been shown to be effective to reduce muscle glycogen content (36, 37).

#### High-intensity interval exercise

The high-intensity interval exercise sessions were preceded by a 5-min warm-up at 90% of the first lactate threshold. Participants completed ten 2-min intervals at an intensity of 20% of the difference between the second lactate threshold and their peak power (182 ± 38 W, 79 ± 5% of peak power). Each 2-min bout was interspersed with a 1-min passive recovery period (33). Participants were required to maintain a pedal frequency of 70-80 rpm during each 2-min bout. The 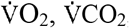, RER, and 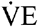 were measured breath-by-breath throughout each high-intensity interval exercise session using the same gas analyzer described for the graded exercise test. Data were then converted to 30-s intervals and the last 30-s of each 2-min bout were used for further analysis.

### Diet and exercise control before starting experimental manipulation

Participants were asked to register all foods and beverages consumed during the 48-h preceding the start of the preliminary test. The usual macronutrient consumption was measured from this diet recall using a computer program designed to perform nutritional calculations (Nutwin, Nutrition Support Program, version 1.5, Department of Health Informatics, Federal University of São Paulo, Brazil). The daily energy and macronutrient intakes during the 48-h preceding the first preliminary test were 3330 ± 475 kcal (43.2 ± 10.0 kcal·kg^-1^), 49% ± 6% carbohydrates (409.3 ± 79.4 g, 5.3 ± 1.4 g·kg^-1^χ 27% ± 5% lipids (101.5 ± 28.2 g, 1.3 ± 0.5 g·kg^-1^), and 24% ± 7% protein (195.0 ± 43.8 g, 2.5 ± 0.5 g·kg^-1^). Participants were asked to replicate the dinner of two days before (day 1), and breakfast, lunch and dinner of one day before the experimental trial (day 2) (Fig. 1). Participants were given verbal and written instructions describing how to repeat this before each subsequent experimental trial. Checklists were used to check any deviations from the menu, and analysis of checklists showed no differences for energy or macronutrient intake between conditions during these meals (Table 1). Participants were also instructed to avoid any strenuous exercise as well as alcohol and caffeine consumption for the 24 h prior to each experimental trial.

**Table 1.** Analysis of checklists for energy or macronutrient intake between conditions for the dinner eaten two days before, and breakfast, lunch and dinner eaten one day before the experimental trial.

|  |  | Dinner two<br>days before the<br>experimental<br>trial | Breakfast one<br>day before the<br>experimental<br>trial | Lunch one<br>day before the<br>experimental<br>trial | Dinner one day<br>before the<br>experimental<br>trial |
| --- | --- | --- | --- | --- | --- |
| **Energy intake**<br>**(Kcal)** | **Control** | 911 ± 348 | 613 ± 358 | 955 ± 233 | 852 ± 215 |
|  | **Twice-a-day** | 918 ± 354 | 602 ± 366 | 925 ± 236 | 886 ± 237 |
|  | **Once-daily** | 887 ± 334 | 602 ± 366 | 948 ± 236 | 884 ± 236 |
| **Carbohydrate**<br>**(%)** | **Control** | 45 ± 19 | 52 ± 12 | 44 ± 5 | 50 ± 9 |
|  | **Twice-a-day** | 45 ± 19 | 53 ± 12 | 46 ± 6 | 49 ± 9 |
|  | **Once-daily** | 43 ± 18 | 53 ± 12 | 44 ± 5 | 49 ± 9 |
| **Protein**<br>**(%)** | **Control** | 28 ± 17 | 17 ± 8 | 32 ± 5 | 21 ± 12 |
|  | **Twice-a-day** | 28 ± 17 | 17 ± 8 | 33 ± 6 | 21 ± 12 |
|  | **Once-daily** | 28 ± 17 | 17 ± 8 | 34 ± 7 | 21 ± 12 |
| **Lipids**<br>**(%)** | **Control** | 27 ± 10 | 31 ± 7 | 23 ± 10 | 29 ± 8 |
|  | **Twice-a-day** | 27 ± 10 | 30 ± 8 | 21 ± 11 | 30 ± 9 |
|  | **Once-daily** | 30 ± 10 | 30 ± 8 | 21 ± 11 | 30 ± 9 |

Data are presented as mean ± standard deviation. n = 8 for all variables.

### Blood collection and analysis

Blood samples were collected from an antecubital vein and separated into three different tubes. Two millilitres of blood were collected in tubes containing sodium fluoride and EDTA (Hemogard™ Fluoride/EDTA, BD Vacutainer®, USA). Blood was centrifuged at 4,000 rev.min^-1^ for 10 min at 4° C with the resulting plasma transferred to 2-mL tubes and immediately analysed for plasma glucose and lactate concentrations. Plasma glucose and lactate concentrations were analysed with a commercially available enzymatic kit (Glucose Liquiform and Enzymatic Lactate, respectively, Labtest, Lagoa Santa, Minas Gerais, Brazil). A further 8 mL of blood was collected in tubes containing Clot activator and gel for serum separation (SST II Plus, BD Vacutainer®, USA), and another 3 mL of blood was collected in tubes containing sodium heparin (Sodium Heparin^N^ Plus, BD Vacutainer®, USA). Both were then centrifuged at 4,000 rev.min^-1^ for 10 min at 4° C, and the resulting serum/plasma frozen and stored in liquid nitrogen for the later analysis of free fatty acid, glycerol, and catecholamine concentrations in the serum/plasma. Serum free fatty acid and glycerol concentrations were determined by an enzymatic colorimetric method (EFFA-100 and EGLY-200, BioAssay, Hayward, California, USA). Plasma catecholamine concentrations were determined by using ion-pairing reverse phase liquid chromatography coupled with electrochemical detection (35).

### Muscle tissue samples and analysis

Nine separate incisions (three per trial) were made into the vastus lateralis under local anaesthesia (2% Xylestesin^®^), and a muscle sample taken using a Bergström needle (38) adapted for manual suction (39). Samples were taken approximately 1 cm apart from a previous biopsy site. Samples [mean: 118 ± 48 mg; range: 49 to 248 mg] were immediately snap-frozen in liquid nitrogen, and then stored at −80°C until subsequent analyses. Muscle samples were taken at rest (pre), immediately after (post) and 3 h after the high-intensity interval exercise (3 h post). Biopsies were subsequently analysed for muscle glycogen content, as well as gene and protein expression (described subsequently).

### Muscle glycogen concentration

Approximately 2 to 3 mg of freeze-dried muscle tissue was powdered and dissected free of all visible non-muscle tissue. Powdered muscle tissue was then extracted with 250 μL of 2 M HCl, incubated at 95°C for 2 h (agitated gently every 20 min), and then neutralized with 750 μL of 0.66 M NaOH. Glycogen concentration was subsequently assayed in triplicate via enzymatic analysis with fluorometric detection (40) and the mean value reported as millimoles per kilogram dry weight.

### Western blotting

#### Muscle homogenate preparations and protein assays

Approximately 20 mg of frozen muscle tissue was homogenized using a TissueLyser II (Qiagen, Valencia, CA) in a 1:20 dilution of ice-cold RIPA buffer (pH 7.4) containing: 0.15 M NaCl, 1% Triton-X100, 0.5% sodium deoxycholate, 0.05 M Tris, 0.1% SDS, 0.1 M EDTA, with the addition of protease/phosphatase inhibitor cocktail (Cell Signalling Technology [CST], #5872, St. Louis, MI). Homogenates were rotated end-over-end for 60 min at 4°C. Protein content of muscle homogenate was measured in triplicate using a Bradford assay (Bio-Rad protein assay dye reagent concentrate, Bio-Rad Laboratories, Hercules, CA) against bovine serum albumin standards (BSA, A9647, Sigma-Aldrich). Nuclear and crude cytosolic fractions were prepared from 40 to 60 mg of wet muscle using a commercially-available nuclear extraction kit (NE-PER^TM^, Pierce, USA). Muscle samples were homogenized in cytoplasmic extraction reagent I buffer containing a protease/phosphatase inhibitor cocktail (CST, 5872). Following centrifugation (16,000 *g* for 5 min at 4° C), the supernatant was taken and pellets containing nuclei were washed five times in PBS to remove cytosolic contamination, before nuclear proteins were extracted by centrifugation (16,000 *g* for 10 min at 4° C) in high-salt nuclear extraction reagent buffer supplemented with the same inhibitors cocktail following manufacturer’s instruction. Verification of subcellular enrichment is presented in Fig. 3 b and 4 b. Sufficient muscle was available to prepare subcellular fractions from all eight participants.

#### Immunoblotting

RIPA-buffered homogenate was diluted in 4X Laemmli buffer (0. 25 M Tris, 8% SDS, 40% glycerol, 0.04% bromophenol blue, 20% 2-mercaptoethanol) and equal amounts of total protein (10 to 20 μg) were loaded on Criterion™ 4-20% TGX Stain-Free™ Precast Gels (Bio-Rad). All samples for a participant were loaded in adjacent lanes on the same gel. Four to six different dilutions of a mixed-homogenate internal standard were also loaded on each gel and a calibration curve plotted of density against protein content. From the subsequent linear regression equation protein abundance was calculated from the measured band intensity for each sample on the gel (41). Gel electrophoresis ran for 90 min at 80-150 V. Proteins were turbo-transferred to a 0.2 μm PVDF membrane at 25 V for 10 min. Membranes were blocked for 60 min at room temperature in 5% non-fat dry milk diluted in Tris-buffered saline with 0.1% Tween-20 (TBST). Membranes were then washed in TBST and incubated overnight at 4°C – with the appropriate primary antibody: Histone H3 (CST, 44995), LDHA (CST, 2012), PGC-1α (CST, 2178), p-ACC^Ser79^ (CST, 3361), AMPK (CST, 2532), p-AMPK^Thr172^ (CST, 2531), p38 MAPK (CST, 9212), p-p38 MAPK^Thr180/Tyr182^ (CST, 9211), PHF20 (CST, 3934), p53 (CST, 2527), TFEB (CST, 9601), and NFAT2 (CST, 8032) diluted (1:1,000) in 5% BSA and 0.02% sodium azide in TBST. Following TBST washes the membranes were incubated in the relevant secondary antibody: Goat anti-rabbit IgG (Perkin Elmer/NEF812001EA), diluted (1:10,000) in 5% non-fat dry milk in TBST, for 60 min at room temperature. After further washes, membranes were developed using Clarity ECL (Bio-Rad) and images were taken with a ChemiDoc Imaging System fitted (Bio-Rad). Densitometry was performed with Image Lab 5.0 software (Bio-Rad). Images are typically displayed with at least five bandwidths above and below the band of interest.

### Real-Time quantitative PCR

#### RNA extraction

Total RNA from approximately 10 to 15 mg of frozen muscle was homogenized in 800 μL of TRIzol reagent (Thermo Fisher Scientific, Waltham, USA) using a TissueLyser II (Qiagen) (42). The concentration and purity of each sample was assessed using a NanoDrop One/One^c^ (Thermo Fisher Scientific). As representative, RNA integrity of a subset of samples was measured using a Bio-Rad Experion microfluidic gel electrophoresis system with Experion RNA StdSens Analysis kit (Bio-Rad, 7007104). All tested samples showed good quality (RNA quality indicator > 7). RNA was stored at −80°C until reverse-transcription was performed.

#### Reverse transcription

1 μg RNA, in a total reaction volume of 20 μL, was reverse-transcribed to cDNA using a Thermocycler (Bio-Rad) and iScript RT Supermix (Bio-Rad, 170-8840) per the manufacturer’s instructions. Priming was performed at 25°C for 5 min and reverse transcription for 30 min at 42°C. All samples, including RT-negative controls, were performed during the same run. cDNA was stored at −20°C until subsequent analysis.

#### qPCR

Relative mRNA expression was measured by qPCR (QuantStudio 7 Flex, Applied Biosystems, Foster City, CA) using SsoAdvanced Universal SYBR Green Supermix (Bio-Rad). Primers were designed using Primer-BLAST (43) to include all splice variants, and were purchased from Sigma-Aldrich (see Supplementary Table 1 for primer details). All reactions were performed in duplicate on 384-well MicroAmp optical plates (4309849, Applied Biosystems) using an epMotion M5073 automated pipetting system (Eppendorf AG). Total reaction volume of 5 μL contained 2 μL of diluted cDNA template, 2.5 μL of mastermix, and 0.3 μM or 0.9 μM primers. All assays ran for 10 min at 95°C, followed by 40 cycles of 15 s at 95°C and 60 s at 60°C. The stability of six potential reference genes were determined by *BestKeeper* (44) and *NormFinder* (45) software, and the three most stably expressed genes were TBP, 18S, and ACTB (Supplementary Table 1). There were no main effects of training approach, time, nor interaction effects, for the quantification cycle (Ct) values of the three most stable housekeeping genes (P > 0.05). Expression of each target gene was normalized to the geometric mean of expression of the three reference genes (46), and using the 2^-ΔΔCt^ method (47).

### Statistical analysis

Statistical analysis was performed using the GraphPad Prism software version 6.01. All data were checked for normality with the Kolomogorov-Smirnov test. To compare the responses before, immediately after, and 3 h after each high-intensity interval exercise session, data were analysed with two-way repeated measures ANOVA (trial vs. time). A Bonferroni post-hoc test was used to locate the differences. All values are expressed as means ± SD. Significance was accepted when p < 0.05.

## RESULTS

### Muscle glycogen concentration

Prior to the high-intensity interval exercise, muscle glycogen concentration was similarly lower in both the twice-a-day and once-daily conditions, compared to the control condition (~45 and 42%, respectively; Fig. 2). Compared to their respective rest values, muscle glycogen concentration decreased similarly after the high-intensity interval exercise in all three conditions (from 245.0 to 135.7 mmol·kg^-1^ dry mass in the twice-a-day, from 262.2 to 163.7 mmol·kg^-1^ dry mass in the once-daily, and from 449.2 to 312.2 mmol·kg^-1^ dry mass in the control; ~45, 38, and 30% of reduction for twice-a-day, once-daily and control, respectively; *P* < 0.05; Fig. 2), and remained lower 3 h post exercise in both the twice-a-day and once-daily conditions, compared to the control condition (~46 and 43%, respectively; Fig. 2).

**Figure 2.**
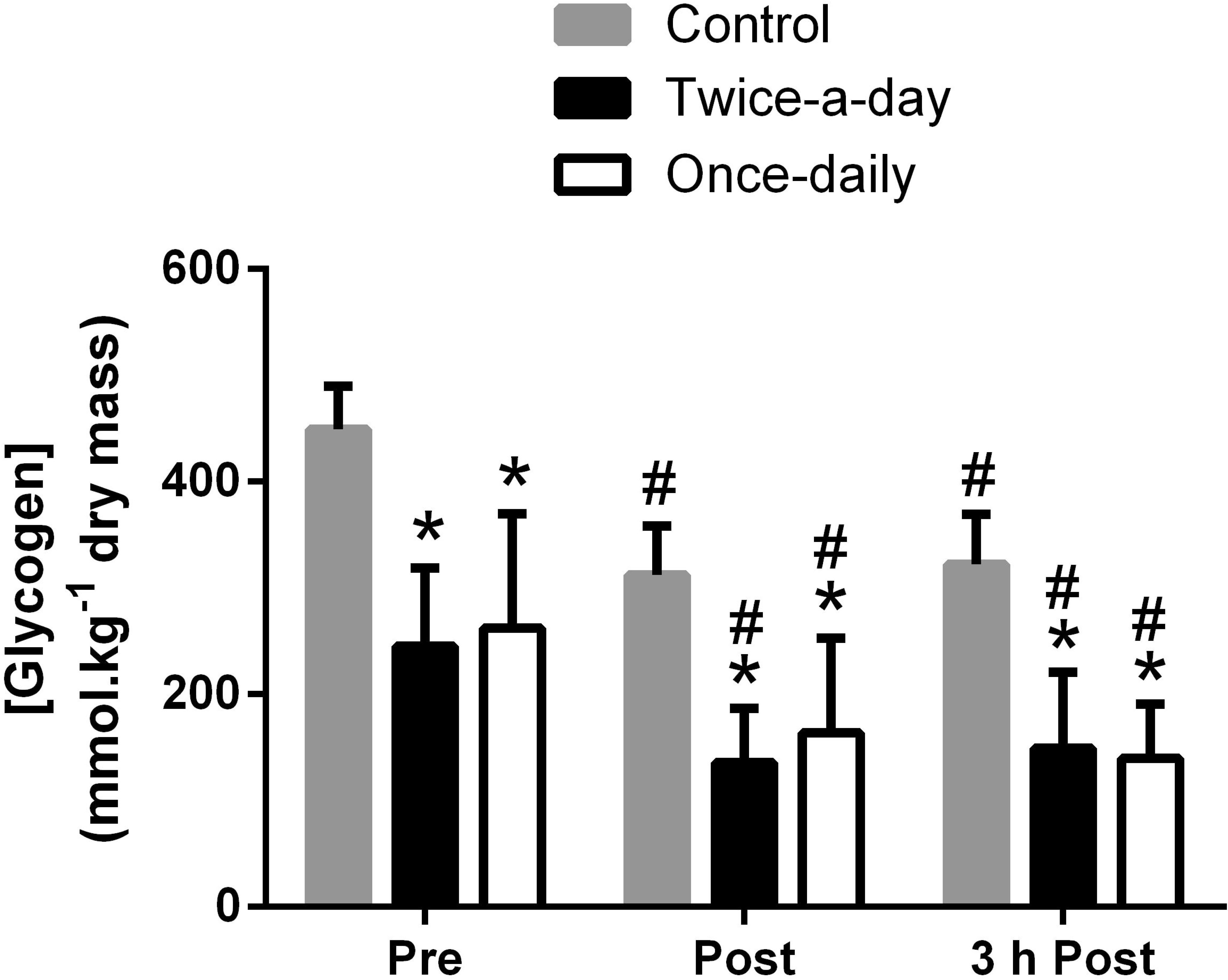
Muscle glycogen concentration pre, post, and 3 h post the high-intensity interval exercise. Data are presented as mean ± standard deviation. n = 8. * significantly lower than control at the same time point (*P* < 0.05); # significantly lower than pre high-intensity interval exercise for the same condition (*P* < 0.05). Two-way analysis of variance (ANOVA) with Bonferroni post hoc test.

### Cytosolic proteins relative abundance

Representative blots are presented in Fig. 3 a, b. Cytosolic p53 protein relative abundance increased immediately post high-intensity interval exercise in all three conditions (Fig. 3 d), with no differences between conditions. Cytosolic PGC-1α, phosphorylated p53 (p-p53^Ser15^), PHF20 protein (PHF20), TFEB, p38 mitogen-activated protein kinase (p38 MAPK), phosphorylated 5’ adenosine monophosphate-activated protein kinase (p-AMPK^Thr172^) and NFAT relative abundance were unaffected by the exercise approach or time (Fig. 3 c, 3 e-j).

**Figure 3.**
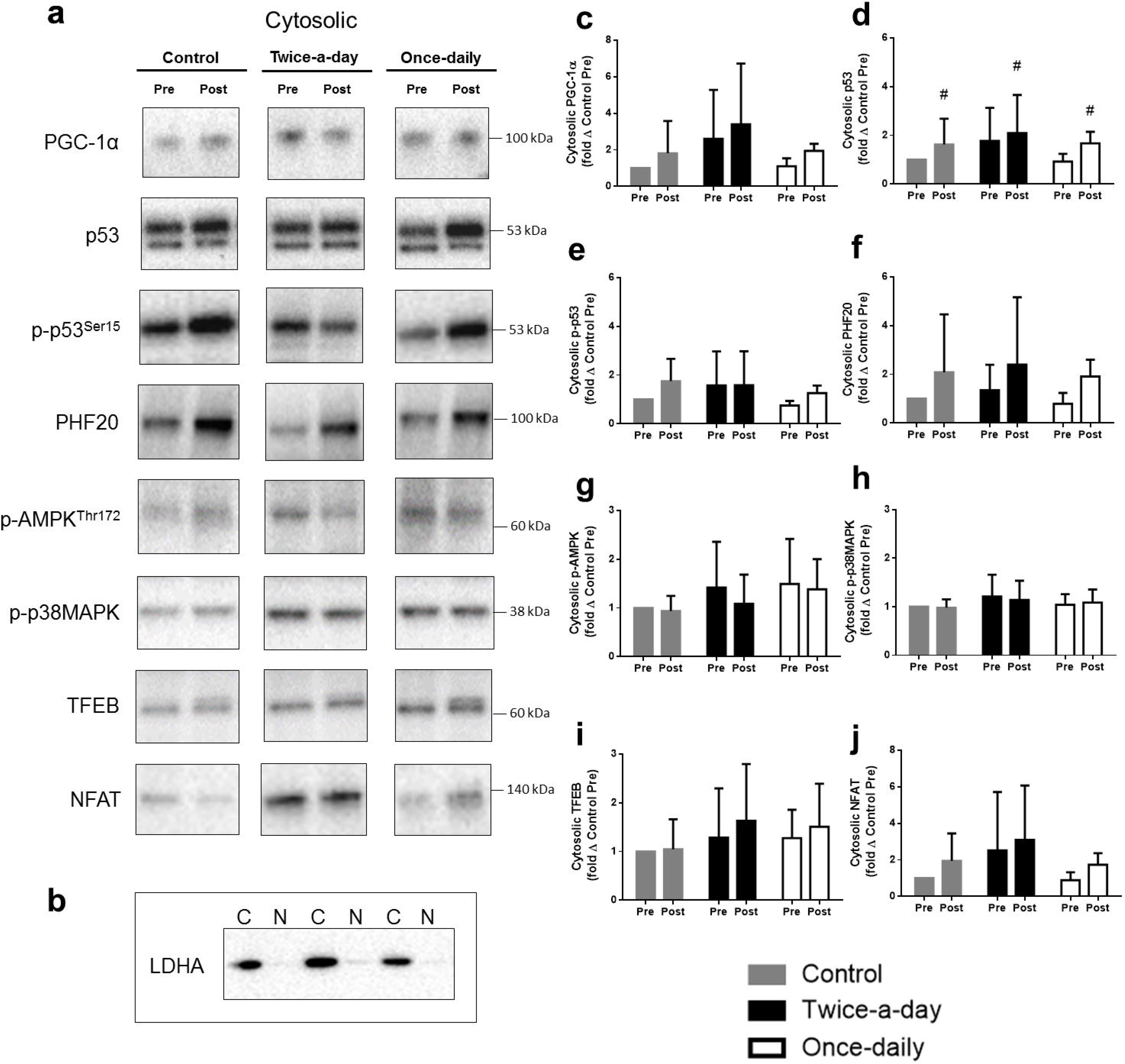
Cytosolic protein relative abundance pre and post the high-intensity interval exercise. **(a)** Representative immunoblots corresponding to total and phosphorylated protein relative abundance measured in the cytosolic fraction, pre and post the high-intensity interval exercise in the control, twice-a-day, and once-daily approaches; **(b)** LDHA was used as indicators of cytosolic enrichment. N: nuclear fractions; C: cytosolic fractions; **(c)** cytosolic peroxisome proliferator-activated receptor-γ coactivator-1 (PGC-1α); **(d)** cytosolic p53 (p53); **(e)** cytosolic phosphorylated p53 (p-p53^Ser15^); **(f)** cytosolic PHF20 (PHF20); **(g)** cytosolic phosphorylated 5’ adenosine monophosphate-activated protein kinase (p-AMPK^Thr172^); **(h)** cytosolic phosphorylated p38 mitogen-activated protein kinase (p-p38MAPK); **(i)** cytosolic transcription elongation factor EB (TFEB); **(j)** cytosolic nuclear factor of activated T cells (NFAT). n = 8 for all proteins. Data are presented as fold changes from control pre (mean ± standard deviation). # significantly higher than pre high-intensity interval exercise for the same condition (*P* < 0.05). Two-way analysis of variance (ANOVA) with Bonferroni post hoc test.

### Nuclear protein relative abundance

Representative blots are presented in Fig. 4 a,b. Nuclear PGC-1α, p53, and p-p53^Ser15^ relative abundance increased post high-intensity interval exercise (Fig. 4 c-e), with no clear differences between the three conditions. The relative abundance of nuclear PHF20, p38MAPK, and p-AMPK^Thr172^ was unaffected by either “train-low” approach or time (Fig. 4 f-h). However, nuclear TFEB relative abundance was greater in the twice-a-day compared to the once-daily condition both pre and post the high-intensity interval exercise (Fig. 4 i). Moreover, nuclear NFAT relative abundance was also greater in the twice-a-day compared to both the once-daily and control condition post the high-intensity interval exercise (Fig. 4 j).

**Figure 4.**
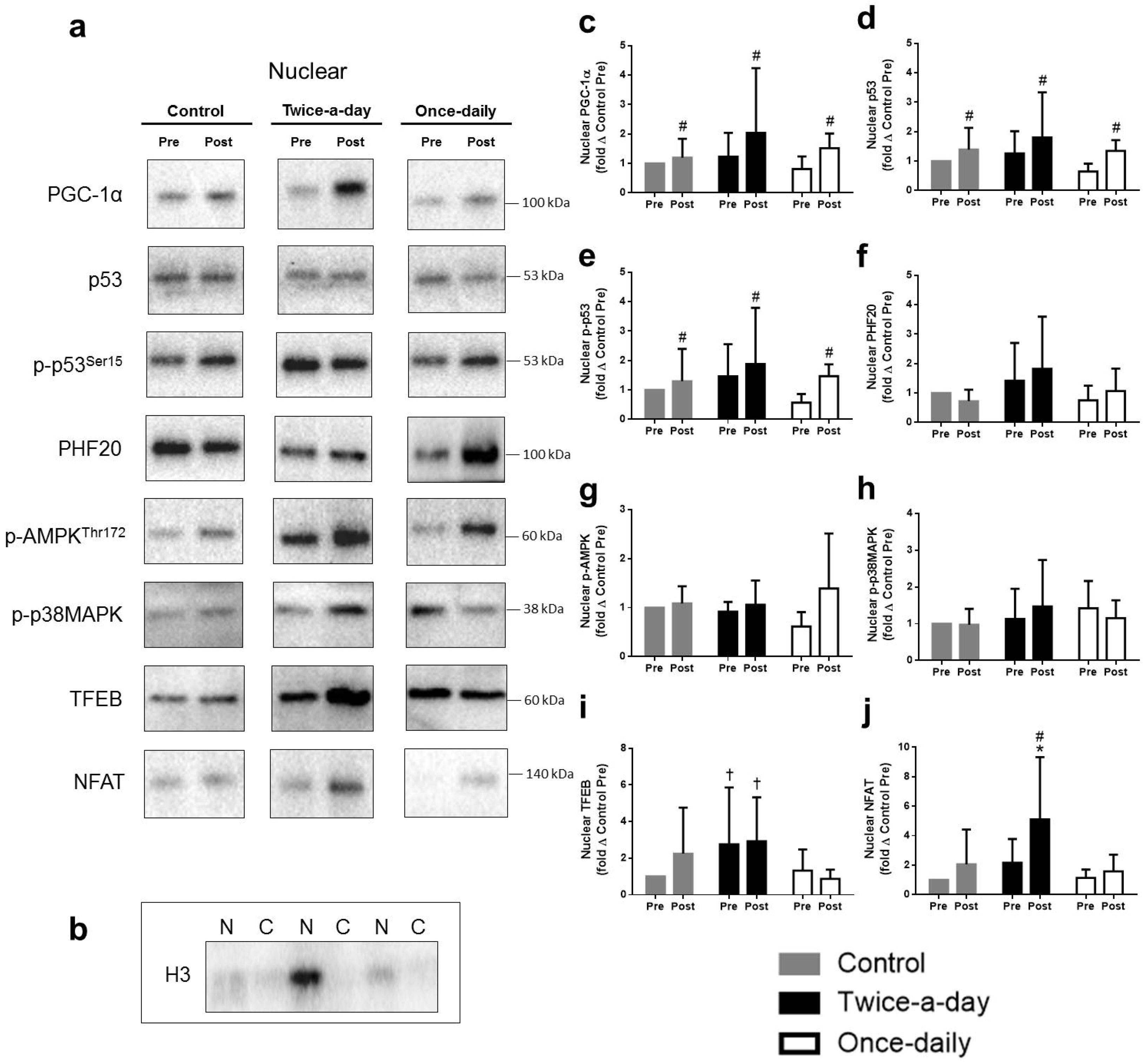
Nuclear protein relative abundance pre and post the high-intensity interval exercise. **(a)** Representative immunoblots corresponding to total and phosphorylated protein relative abundance measured in the nuclear fraction, pre and post the high-intensity interval exercise in the control, twice-a-day, and once-daily approaches; **(b)** H3 was used as indicators of nuclear enrichment. N: nuclear fractions; C: cytosolic fractions; **(c)** nuclear peroxisome proliferator-activated receptor-γ coactivator-l (PGC-1α); **(d)** nuclear p53 (p53); **(e)** nuclear phosphorylated p53 (p-p53^Ser15^); **(f)** nuclear PHF20 (PHF20); **(g)** nuclear phosphorylated AMPKThr172 (p-AMPK^Thr172^); (h) nuclear phosphorylated p38MAPK (p-p38MAPK); **(i)** nuclear transcription elongation factor EB (TFEB); **(j)** nuclear factor of activated T cells (NFAT). n = 8 for all proteins. Data are presented as fold changes from control pre (mean ± standard deviation). * significantly higher than the once-daily and control condition at the same time point (*P* < 0.05); † significantly higher than the once-daily condition at the same time point (*P* < 0.05). # significantly higher than pre high-intensity interval exercise for the same condition (*P* < 0.05). Two-way analysis of variance (ANOVA) with Bonferroni post hoc test.

### Mitochondrial-related gene expression

Pre high-intensity interval exercise, total PGC-1α mRNA content was ~9-fold higher in the twice-a-day compared to both the once-daily and control conditions (Fig. 5 a). Three hours post the high-intensity interval exercise, total PGC-1α mRNA content increased in all three conditions compared with their respective pre-values; however, total PGC-1α mRNA content remained ~ 10-fold higher in the twice-a-day compared to both control and once-daily conditions. Similarly, PGC-1α isoform 4 mRNA content was ~24-fold higher at pre high-intensity interval exercise in the twice-a-day compared to both the once-daily and control conditions (Fig. 5 c). At 3 h post high-intensity interval exercise, the PGC-1α isoform 4 mRNA content increased in all three conditions compared with their respective pre-values; however, the PGC-1α isoform 4 mRNA content remained ~10-fold higher in the twice-a-day compared to the control and once-daily conditions. Additionally, PGC-1α isoform 1 mRNA content was higher in the twice-a-day compared to the once-daily when all time points where considered (Fig. 5 b). There was no effect of condition for p53, TFEB, chromodomain-helicase-DNA-binding protein 4 (CHCHD4), p21, mitochondrial transcription factor A (Tfam), NADH dehydrogenase subunit β (NDUFβ; mitochondrial complex I), succinate dehydrogenase subunit β (SDHβ; mitochondrial complex II), cytochrome c (mitochondrial complex III), and cytochrome c oxidase subunit IV (COX IV; mitochondrial complex IV) mRNA content post high-intensity interval exercise (Fig. 5 d-l). However, the mRNA content of representative subunits of mitochondrial complexes II, III, and IV increased 3 h post high-intensity interval exercise to a similar extent for all three conditions.

**Figure 5.**
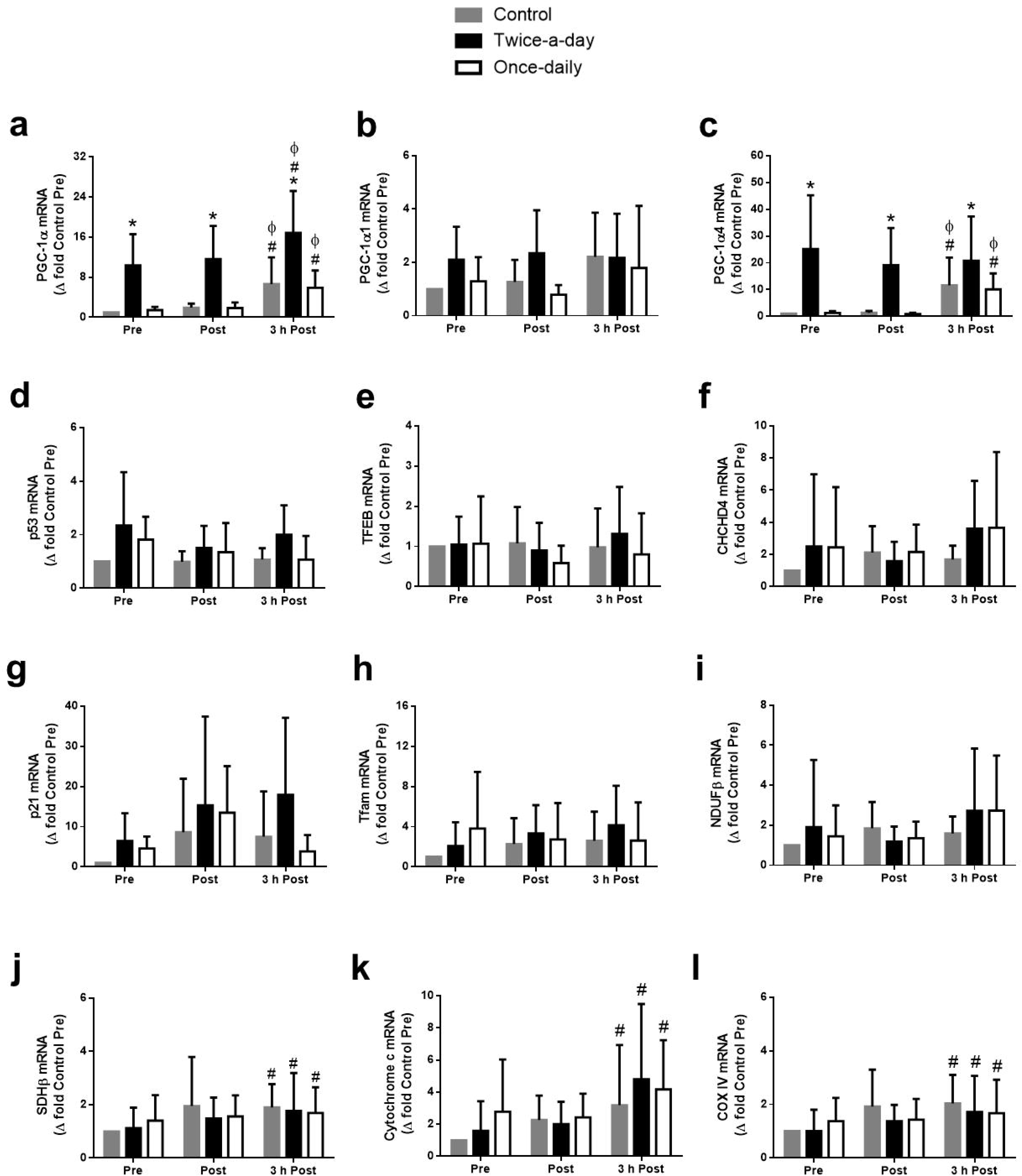
Mitochondrial-related gene expression pre, post, and 3 h post the high-intensity interval exercise. **(a)** peroxisome proliferator-activated receptor-coactivator-1 (PGC-1α) total gene expression; **(b)** PGC-1α isoform 1 gene expression; **(c)** PGC-1α isoform 4 gene expression; **(d)** p53 (p53) gene expression; **(e)** transcription elongation factor EB (TFEB) gene expression; **(f)** Chromodomain-helicase-DNA-binding protein 4 (CHCHD4) gene expression; **(g)** p21 protein (p21) gene expression; **(h)** mitochondrial transcription factor A (Tfam) gene expression; (i) NADH dehydrogenase (NDUFβ) (mitochondrial complex I) gene expression; **(j)** succinate dehydrogenase subunit β (SDHβ) (mitochondrial complex II) gene expression; **(k)** cytochrome c (mitochondrial complex III) gene expression; (l) cytochrome c oxidase subunit IV (COXIV) (mitochondrial complex III) gene expression. n = 8 for all genes (except Tfam, n = 7). Data are presented as fold changes from control pre (mean ± standard deviation). * significantly higher than the once-daily and control condition at the same time point (*P* < 0.05); # significantly higher than pre high-intensity interval exercise for the same condition (*P* < 0.05); □ significantly higher than post high-intensity interval exercise for the same condition (*P* < 0.05). Two-way analysis of variance (ANOVA) with Bonferroni post hoc test.

### Fat transport and lipolysis related genes

The content of carnitine palmitoyltransferase I subunit A (CPT1A) mRNA was higher in both the twice-a-day and once-daily conditions compared with the control condition at 3 h post the high-intensity interval exercise; however, there was no difference between the two “train-low” approaches (Fig. 6 a). There was an increase in mitochondrial uncoupling protein 3 (UCP3) mRNA content 3 h post exercise in all three conditions. The UCP3 mRNA content was, however, significantly higher only in the twice-a-day compared to the control condition at 3 h post the high-intensity interval exercise (Fig. 6 b). Pre high-intensity interval exercise, the PPARα mRNA content was ~11-fold higher in the twice-a-day compared to both the once-daily and control conditions (Fig. 6 c). Three hours post the high-intensity interval exercise, the PPARα mRNA content increased ~7- and 9-fold in the once-daily and control conditions, respectively, compared with their respective pre-values; however, the PPARα mRNA content remained ~16-fold higher in the twice-a-day compared to the once-daily and control conditions. The PPARβ/δ mRNA content was higher in the twice-a-day than in the control condition post the high-intensity interval exercise, and higher than both the once-daily and control conditions at 3 h post the high-intensity interval exercise (Fig. 6 d). The PPARβ/δ mRNA content was higher at 3 h post the high-intensity interval exercise compared to pre- and post-only for the twice-a-day approach. The citrate synthase (CS) mRNA content was higher 3 h post the high-intensity interval exercise compared to post for all three conditions. However, the peroxisome proliferator-activated receptor gamma (PPARγ), β-hydroxyacyl-CoA dehydrogenase (β-HAD), and fatty acid translocase cluster of differentiation 36 (CD-36) mRNA content were unaffected by the “train-low” approach or time (Fig. 6 e-h).

**Figure 6.**
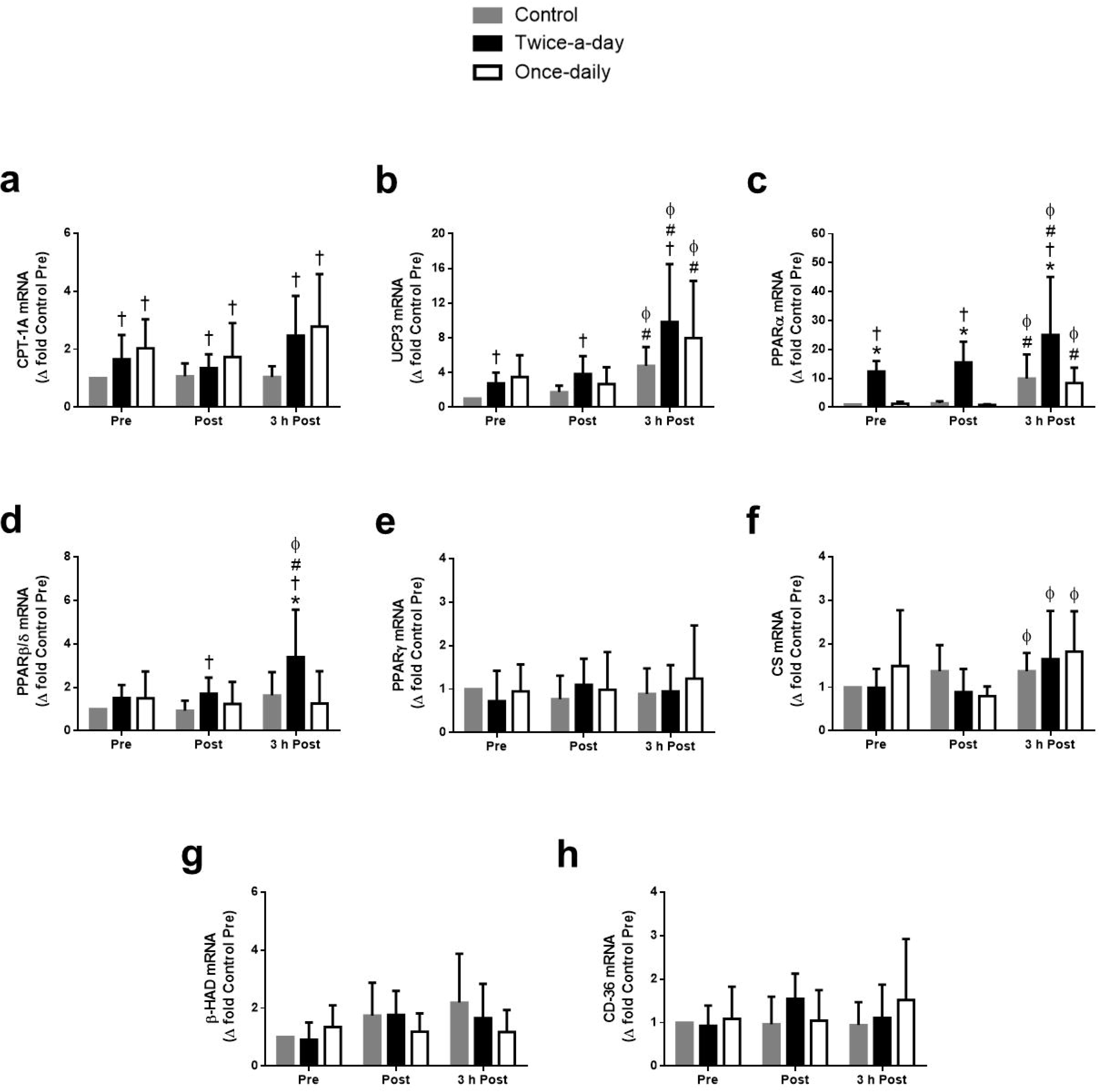
Genes related to fat transport and lipolysis pre, post, and 3 h post the high-intensity interval exercise. **(a)** carnitine palmitoyltransferase I (CPT1) gene expression; **(b)** mitochondrial uncoupling protein 3 (UCP3) gene expression; **(c)** peroxisome proliferator-activated receptor (PPARα) gene expression; **(d)** peroxisome proliferator-activated receptor β/δ (PPARβ/δ) gene expression; **(e)** peroxisome proliferator-activated receptor γ (PPARγ) gene expression; **(f)** citrate synthase (CS) gene expression; **(g)** β-hydroxyacyl-CoA dehydrogenase (β-HAD) gene expression; **(h)** fatty acid translocase cluster of differentiation 36 (CD-36) gene expression. n = 8 for all genes (except CS and UCP3, where n = 7). Data are presented as fold changes from control pre (mean ± standard deviation). * significantly higher than the once-daily condition at the same time point (*P* < 0.05); † significantly higher than the control condition at the same time point (*P* < 0.05); # significantly higher than pre high-intensity interval exercise for the same condition (*P* < 0.05); □ significantly higher than post high-intensity interval exercise for the same condition (*P* < 0.05). Two-way analysis of variance (ANOVA) with Bonferroni post hoc test.

### Glycolysis related genes

The mRNA content of phosphofructokinase (PFK) and glucose transporter 4 (GLUT4) was unaffected by the exercise approach or time. While pyruvate dehydrogenase kinase isoenzyme 4 (PDK4) mRNA content increased 3 h post the high-intensity interval exercise, there was no difference for the three different exercise approaches (Fig. 7 a-c).

**Figure 7.**
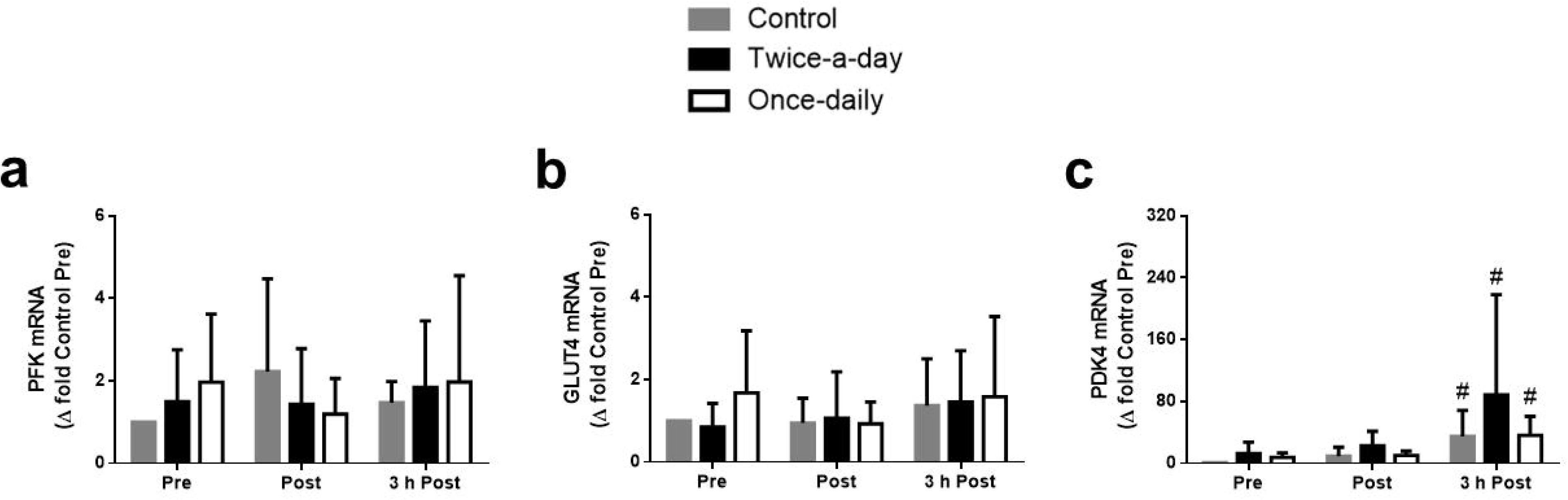
Genes related to carbohydrate metabolism pre, post, and 3 h post the high-intensity interval exercise. **(a)** phosphofructokinase (PFK) gene expression; **(b)** glucose transporter 4 (GLUT4) gene expression; **(c)** pyruvate dehydrogenase kinase isoenzyme 4 (PDK4) gene expression. n = 8 for all genes. Data presented as fold changes from control pre (mean ± standard deviation). # significantly higher than pre high-intensity interval exercise for the same condition (*P* < 0.05). Two-way analysis of variance (ANOVA) with Bonferroni post hoc test.

### Physiological responses

The twice-a-day approach was associated with a higher heart rate, ventilation, and oxygen uptake, and a lower plasma glucose concentration, during high-intensity interval exercise than both the once-daily and the control condition (Fig. 8 a-c and Table 2). Plasma glucose was also lower during the high-intensity interval exercise in the once-daily compared to the control condition (Table 2). In addition, the respiratory exchange ratio was lower and the serum free fatty acid concentration higher than the control only for the twice-a-day approach (Fig. 8 d and Table 2). Serum glycerol was higher and plasma lactate lower post the high-intensity interval exercise in both the twice-a-day and once-daily conditions compared with the control condition (Table 2). Plasma epinephrine and norepinephrine concentrations were not influenced by the exercise approach undertaken.

**Figure 8.**
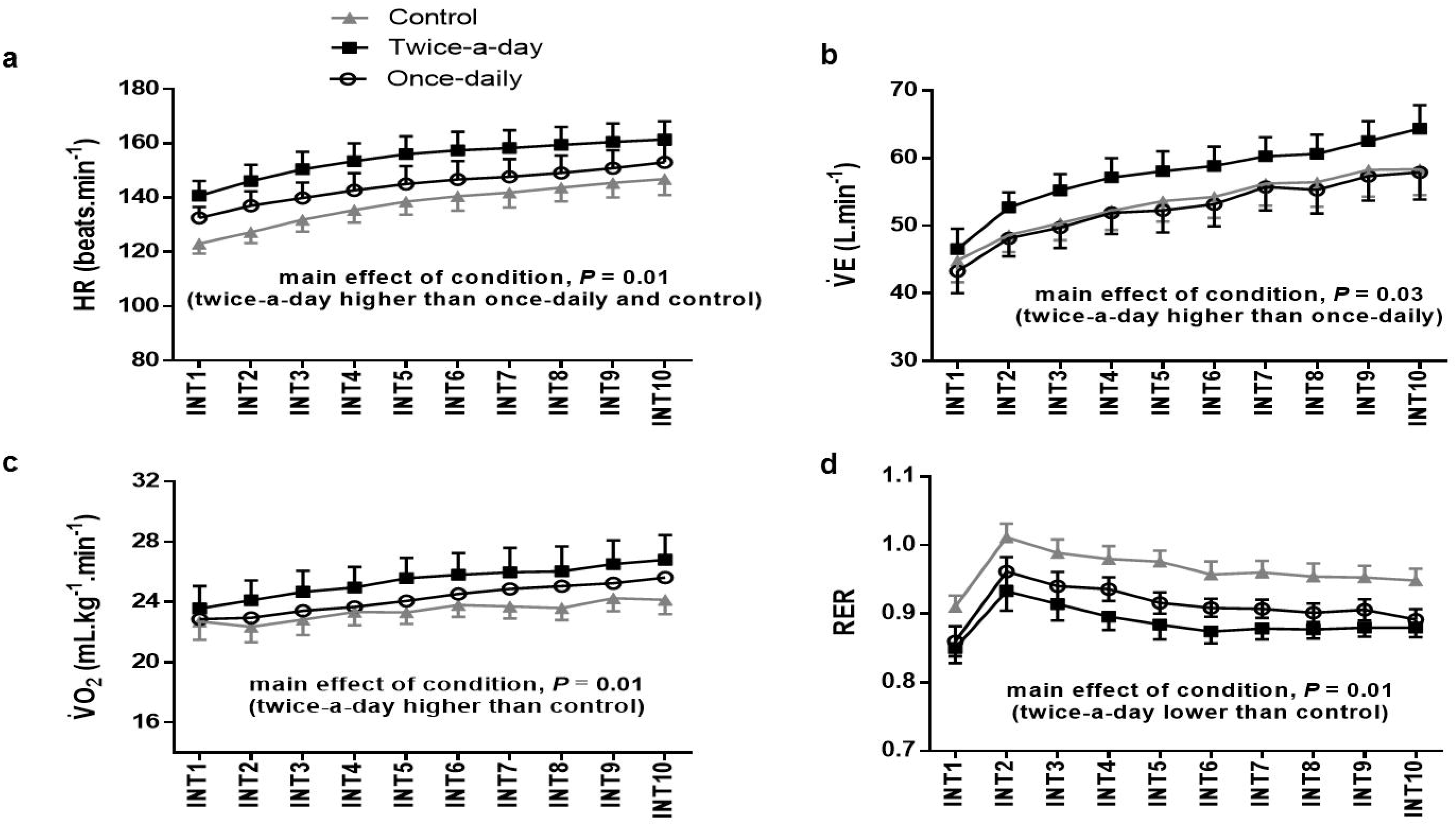
Physiological and systemic responses during the high-intensity interval exercise. **(a)** Heart rate (HR); **(b)** Ventilation 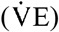; **(c)** Oxygen uptake 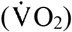; **(d)** Respiratory exchange ratio (RER). n = 8 for all variables. Data are presented as mean ± standard deviation. Time effect has been omitted for clarity. INT: Interval. Two-way analysis of variance (ANOVA) with Bonferroni post hoc test.

**Table 2.**
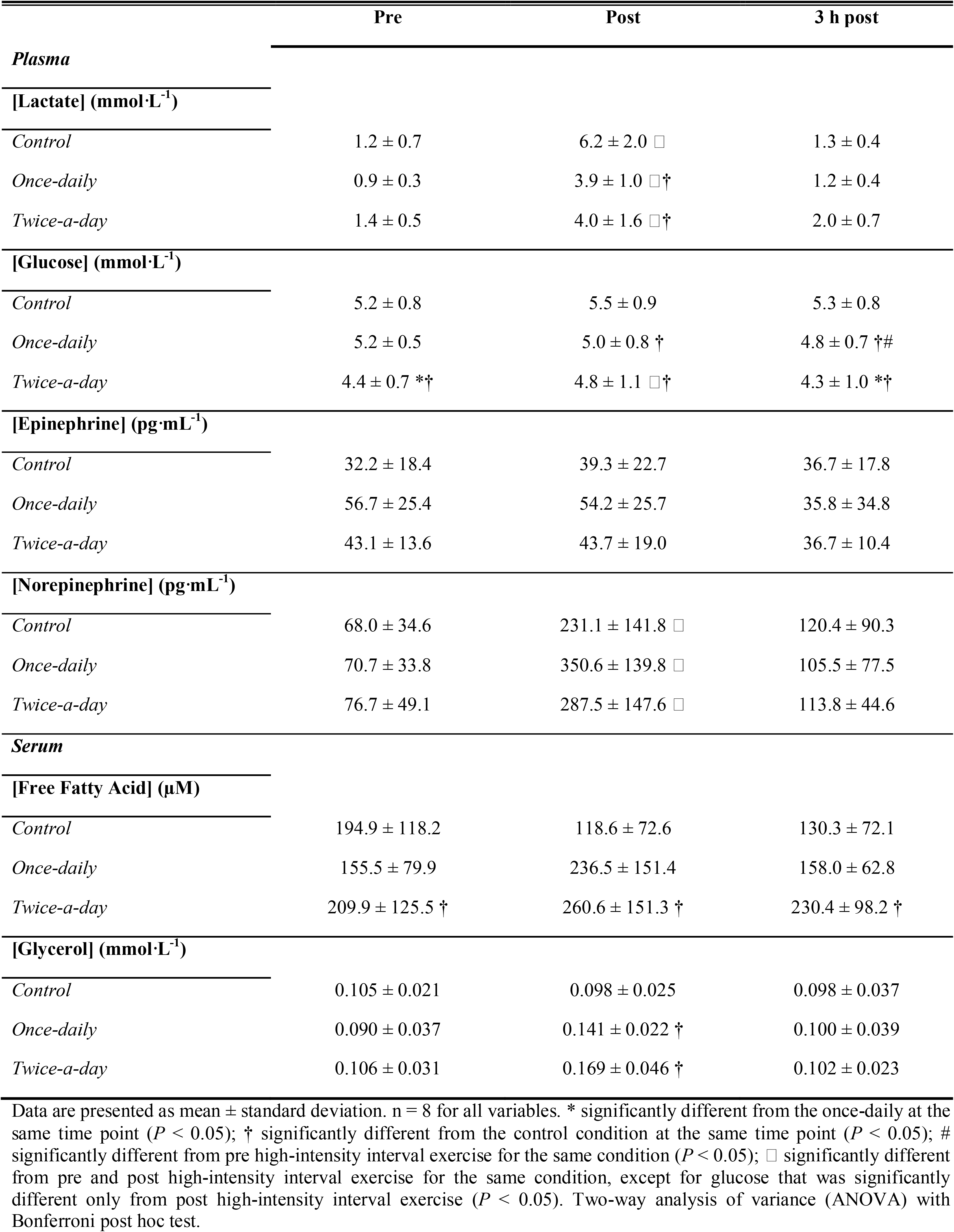
Plasma lactate, glucose, epinephrine and norepinephrine concentrations, and serum free fatty acid and glycerol concentrations pre, post and 3 h post the high-intensity interval exercise.

|  | Pre | Post | 3 h post |
| --- | --- | --- | --- |
| **Plasma** |  |  |  |
| **[Lactate] (mmol·L<sup>-1</sup>)** |  |  |  |
| *Control* | 1.2 ± 0.7 | 6.2 ± 2.0 □ | 1.3 ± 0.4 |
| *Once-daily* | 0.9 ± 0.3 | 3.9 ± 1.0 □ † | 1.2 ± 0.4 |
| *Twice-a-day* | 1.4 ± 0.5 | 4.0 ± 1.6 □ † | 2.0 ± 0.7 |
| **[Glucose] (mmol·L<sup>-1</sup>)** |  |  |  |
| *Control* | 5.2 ± 0.8 | 5.5 ± 0.9 | 5.3 ± 0.8 |
| *Once-daily* | 5.2 ± 0.5 | 5.0 ± 0.8 † | 4.8 ± 0.7 † # |
| *Twice-a-day* | 4.4 ± 0.7 * † | 4.8 ± 1.1 □ † | 4.3 ± 1.0 * † |
| **[Epinephrine] (pg·mL<sup>-1</sup>)** |  |  |  |
| *Control* | 32.2 ± 18.4 | 39.3 ± 22.7 | 36.7 ± 17.8 |
| *Once-daily* | 56.7 ± 25.4 | 54.2 ± 25.7 | 35.8 ± 34.8 |
| *Twice-a-day* | 43.1 ± 13.6 | 43.7 ± 19.0 | 36.7 ± 10.4 |
| **[Norepinephrine] (pg·mL<sup>-1</sup>)** |  |  |  |
| *Control* | 68.0 ± 34.6 | 231.1 ± 141.8 □ | 120.4 ± 90.3 |
| *Once-daily* | 70.7 ± 33.8 | 350.6 ± 139.8 □ | 105.5 ± 77.5 |
| *Twice-a-day* | 76.7 ± 49.1 | 287.5 ± 147.6 □ | 113.8 ± 44.6 |
| **Serum** |  |  |  |
| **[Free Fatty Acid] (μM)** |  |  |  |
| *Control* | 194.9 ± 118.2 | 118.6 ± 72.6 | 130.3 ± 72.1 |
| *Once-daily* | 155.5 ± 79.9 | 236.5 ± 151.4 | 158.0 ± 62.8 |
| *Twice-a-day* | 209.9 ± 125.5 † | 260.6 ± 151.3 † | 230.4 ± 98.2 † |
| **[Glycerol] (mmol·L<sup>-1</sup>)** |  |  |  |
| *Control* | 0.105 ± 0.021 | 0.098 ± 0.025 | 0.098 ± 0.037 |
| *Once-daily* | 0.090 ± 0.037 | 0.141 ± 0.022 † | 0.100 ± 0.039 |
| *Twice-a-day* | 0.106 ± 0.031 | 0.169 ± 0.046 † | 0.102 ± 0.023 |

Data are presented as mean ± standard deviation. n = 8 for all variables. \* significantly different from the once-daily at the same time point ( $P < 0.05$ ); † significantly different from the control condition at the same time point ( $P < 0.05$ ); # significantly different from pre high-intensity interval exercise for the same condition ( $P < 0.05$ ); □ significantly different from pre and post high-intensity interval exercise for the same condition, except for glucose that was significantly different only from post high-intensity interval exercise ( $P < 0.05$ ). Two-way analysis of variance (ANOVA) with Bonferroni post hoc test.

## DISCUSSION

There is continued debate about whether beginning exercise with low muscle glycogen stores potentiates the exercise-induced increase in genes associated with mitochondrial biogenesis and metabolism (10, 11, 48–52). Some of this controversy may relate to the observation that some of the evidence supporting the “train-low” approach is based on performing the experimental exercise session a few hours after a glycogen-lowering exercise session (20–23, 25–27). Thus, it is difficult to determine if any observed effects are due to performing the second exercise session with low muscle glycogen stores and/or performing the second exercise session close to the first. We investigated this question by performing the same exercise session with similar starting muscle glycogen stores, but either 2 or 15 h following the previous glycogen-lowering exercise session. Our results indicate that the greater exercise-induced nuclear protein abundance (TFEB and NFAT) and transcription of genes involved in mitochondrial biogenesis (PGC-1α, PPARα, PPARβ/δ) with the so-called “train-low” approach might be attributed to performing two exercise sessions in close proximity.

Despite the different recovery periods between exercise sessions for the two “train-low” approaches, muscle glycogen prior to the high-intensity interval exercise was reduced to a similar extent in both the twice-a-day and once-daily condition compared with the control condition (Fig. 2). This level of muscle glycogen concentration is consistent with previous studies that have utilized either the twice-a-day (21, 22) or once-daily (19) approach. In addition, muscle glycogen concentration before commencing the high-intensity exercise was below 300 mmol·kg^-1^dry mass for both the twice-a-day and once-daily conditions. Post high-intensity interval exercise muscle glycogen concentration remained above 100 mmol·kg^-1^dry mass. These values have been suggested as an “upper” and “lower” limit threshold, respectively, for which muscle glycogen levels may modulate genes related to mitochondrial biogenesis (52). Thus, the similar low muscle glycogen levels when commencing the high-intensity interval exercise in both “train-low” conditions allowed us to determine whether any differences for exercise-induced gene or protein expression could be attributed to performing high-intensity interval exercise with reduced muscle glycogen stores or to performing high-intensity interval exercise close to a previous exercise session.

In the present study, both p-AMPK and p-p38 MAPK relative protein abundance in both the cytosolic and nucleus fractions was not altered with exercise or influenced by either “train-low” approach (Fig. 4 g,h). It has been suggested that the greater exercise-induced upregulation of cell-signalling pathways with reduced muscle glycogen content may be associated with enhanced nuclear and cytosolic abundance and greater activation of p-AMPK and p-p38 MAPK (14, 15, 53), but contradictory results have been reported (19, 21, 25, 54). For example, Cochran et al. (25) observed larger increases in cytosolic p-p38 MAPK but not p-AMPK with the twice-a-day approach, while Yeo et al. (21) reported larger increases in cytosolic p-AMPK but without alterations in p-p38 MAPK with the twice-a-day compared to the once-daily approach. On the other hand, Gejl et al. (54) and Psilander et al. (19) found no effect of either the twice-a-day or once-daily approach on both p-AMPK and p-p38 MAPK protein content in cytosolic fractions. In addition, Chan et al. (14) and Steinberg et al. (15) reported an increase in nuclear abundance of p-p38 MAPK and AMPK, respectively, with the once-daily approach. The explanation for these contradictory results is unclear, but it is worth noting both the glycogen-depleting exercise and the “depleted-exercise” varied considerably in these studies; either high- (25, 54) or low-intensity (14, 15, 19, 21) exercise was used to reduce muscle glycogen, and high- (21, 25) or low-intensity (14, 15, 19, 54) exercise was implemented during the second exercise session (depleted-exercise). However, activation of AMPK and p38 MAPK does not seem to be associated with muscle glycogen levels or a particular train-low regime (55). Thus, it remains to be solved which protocol or strategy might induce augmented activation of p-AMPK and p-p38 MAPK.

Although p-AMPK and p-p38 MAPK relative protein abundance in the cytosolic and nucleus were not altered with exercise or “train-low” approaches, there was greater PGC-1α, p-p53, and p53 relative protein abundance in the nucleus immediately post the high-intensity interval exercise (Fig. 4 c-e). There was also a higher p53 cytosolic protein abundance post the high-intensity interval exercise (Fig 3 d). The increased nuclear and cytosolic p53 abundance are consistent with the well-accepted notion that cellular stress is associated with accumulation of p53 protein - a process that has been partly attributed to increased p53 protein stability (56, 57). Nevertheless, we found no significant differences between either of the “train-low” approaches and the control condition for p-p53 and p53 relative protein abundance in either subcellular fraction. The “train-low” approach has been associated with a greater exercise-induced cytosolic relative protein abundance of p-p53 (16), which is suggested to influence the gene expression of PGC-1α (58). As differences in neither p-p53 nor p53 protein content in the nucleus could explain the greater exercise-induced increases in PGC-1α gene expression with the different train-low approaches in the present study, we subsequently investigated the nuclear abundance of other proteins that may contribute to the exercise-induced regulation of this gene.

The nuclear abundance of NFAT increased significantly after performing high-intensity interval exercise only in the twice-a-day condition, and post-exercise values were higher than both the control and once-daily conditions (Fig. 4 j). In addition, in both the pre- and post-exercise muscle samples the nuclear abundance of TFEB was significantly greater with the twice-a-day compared to the once-daily approach (Fig. 4 i). As activated calcineurin dephosphorylates both NFAT (59, 60) and TFEB (61–63), leading to their translocation to the nucleus (59–64), this suggests there might be greater calcineurin activation when high-intensity interval exercise is performed soon after a prior exercise session. Although we did not have sufficient muscle sample to test this hypothesis, in the twice-a-day condition we observed significantly greater exercise-induced increases of genes that have been reported to be regulated by calcineurin (e.g., PGC-1α, PPARα and PPARβ/δ; (64, 65)).

In contrast to our findings, some studies have reported greater exercise-induced increases in PGC-1α mRNA with the once-daily approach (~6-fold from rest) when compared to a control condition (~3-fold from rest) (16, 18, 19). This previously reported ~6-fold increase in PGC-1α mRNA with the once-daily approach is of similar magnitude to the PGC-1α mRNA increase with the once-daily approach in the present study (~6 fold from rest), although this was not significantly different to the change we observed in the control condition (~6-fold from rest). However, we did observe a significantly greater exercise-induced increase in PGC-1α (and PPARα, PPARβ/δ) mRNA content in the twice-a-day condition, even though muscle glycogen content before commencing the high-intensity interval exercise was similar between twice-a-day and once-daily conditions, and both were lower than control. Our findings therefore suggest that the greater exercise-induced increase in these genes in our study can be attributed to performing the high-intensity interval exercise soon after the prior exercise session, rather than beginning the high-intensity interval exercise with lowered muscle glycogen levels. It should be mentioned, however, that carbohydrate might still play a role in myocellular signalling as ingesting exogenous carbohydrate between the two exercise sessions has been reported to block the stimulus for inducing oxidative enzyme adaptations in skeletal muscle (25, 27).

In the twice-a-day condition we also observed a greater circulating free fatty acid concentration pre, post, and 3 h post the high-intensity interval exercise (Table 2). Although respiratory exchange ratio values were always below 1.0 after the third interval, and systematically lower in the twice-a-day compared to the control condition (Fig. 8), which could be indicative of greater fat oxidation, this conclusion is limited by the use of nonprotein respiratory quotient to calculate substrate oxidation during nonsteady state exercise (66). The increase in circulating free fatty acids, however, might activate calcineurin (67) and regulate skeletal muscle metabolism via coordinated changes in gene expression (64). As previously mentioned, activated calcineurin will lead to the translocation of both NFAT and TFEB to the nucleus (59–64) and the subsequent upregulation of genes related to fat metabolism (64). These exercise-induced changes are consistent with studies that have reported greater increases in fat oxidation when training twice-a-day (20, 22). Thus, these results reinforce the hypothesis that twice-a-day training might be more effective to induce adaptions related to fat metabolism.

Our study has some limitations that should be mentioned. Firstly, as we were constrained by the number of muscle biopsies we could take, we elected to take our final muscle biopsy 3 h post exercise as most of the key genes investigated in the present study have been reported to peak ~3 h after exercise (e.g., PGC-1α and PPARβ/δ) (5, 68). However, other genes might peak later (e.g. CD-36) (13) and we may have underestimated the influence of the two “train-low” approaches on some genes.

Similarly, we did not perform muscle biopsies immediately and 3 h after the muscle glycogen-depleting exercise; therefore, relevant changes in molecular signalling following the first exercise session in the once-daily condition may have been missed. Secondly, consistent with most related research (16, 18, 23, 25–27), we recruited participants who were active (~3.3 h of aerobic training per week) but who were not well-trained athletes. While this may limit the applicability of our findings to elite athletes, it is difficult to perform similar experiments requiring multiple muscle biopsies in well-trained athletes. Thus, further studies comparing different “train-low” approached in well-trained athletes are required.

In summary, our findings indicate that the greater exercise-induced signalling with the so-called “train-low” approach in our study can partially be attributed to the performance of two exercise sessions in close succession rather than exercising with a reduced muscle glycogen content. We presented evidence that performing two exercise sessions separated by a short recovery period increases the nuclear abundance of TFEB and NFAT and potentiates the transcription of PGC-1α, PPARα and PPARβ/δ. Although we have proposed some molecular mechanisms by which the twice-a-day approach might be a more effective strategy to induce adaptations related to mitochondrial biogenesis and fat oxidation, further research is required to determine if training using the twice-a-day approach results in greater changes in mitochondrial content, mitochondrial function, and fat oxidation.

## Acknowledgements

This work was supported by the Brazilian National Council for Scientific and Technological Development (Special visiting researcher of science without borders program, process N° 406201/2013-7) and an Australian Research Council Grant (DP140104165) to D. J. Bishop. V.A. Andrade-Souza is grateful to the Foundation for the Support of Science and Technology of the State of Pernambuco (FACEPE) for his scholarship. We thank G. McConnel, X. Yan, C. Granata, A.J. Genders, E.C. Marin, R. Aragão and C.R. Correia-Oliveira for technical assistance.

## Author contributions

V. A. A-S., R. B., D. J. B., and A. E. L-S. designed the experiments. V. A. A-S., T. G., A. S., K. A. S. S., F. T., L. A., and A. E. L-S. performed the experiments. V. A. A-S., T. G., A. S., K. A. S. S., F. T., L. A., and A. E. L-S. conducted the exercise analyses at Department of Physical Education and Sports Science, Academic Center of Vitoria, Federal University of Pernambuco and V. A. A-S., J. F., E. P., N. S., and J. K. conducted qPCR and western blot analyses at Institute for Health and Sport, Victoria University. V. A. A-S analyzed data and discussed analyses and results with J. F., J. K., D. J. B., and A. E. L-S. T.G., K. A. S. S., R. B., and C. G. L. supported data analyses. V. A. A-S., K. A. S. S., J. K., D. J. B., and A. E. L-S made the figures. All authors wrote and approved the manuscript.

## Nonstandard Abbreviations

TFEB: transcription factor EB;
NAFT: nuclear factor of activated T cells;
PGC-lα: peroxisome proliferator-activated receptor-□ coactivator l alpha;
PPARα: peroxisome proliferator-activated receptor alpha;
PPARβ/δ: peroxisome proliferator-activated receptor beta/delta;
CHO: carbohydrate;
p53: p53 protein;
PHF20: PHF20 protein;
p38MAPK: p38 mitogen-activated protein kinase;
AMPK: 5’ adenosine monophosphate-activated protein kinase;
COX IV: cytochrome c oxidase subunit IV;
CPT1: carnitine palmitoyltransferase 1;
NDUF: NADH: ubiquinone oxidoreductase;
SDH: succinate dehydrogenase;
GLUT4: Glucose transporter type 4;
β-HAD: 3-hydroxyacyl-CoA dehydrogenase;
CD36: fatty acid translocase cluster of differentiation 36;
PFK: phosphofructokinase;
CS: citrate synthase;
PPARα: peroxisome proliferator-activated receptor alpha;
PPAR□: peroxisome proliferator-activated receptor delta;
UCP3: uncoupling protein 3;
Tfam: mitochondrial transcription factor A;
PDK4: pyruvate dehydrogenase kinase 4;
CHCHD4: coiled-coil-helix-coiled-coil-helix domain containing 4;
p21: p21 protein;
GAPDH: glyceraldehyde 3-phosphate dehydrogenase;
B2M: β-2-microglobulin;
TBP: TATA-box binding protein;
18S: 18S ribosomal RNA;
ACTB: actin beta.

