## Supplementary Table 1 for "Exercise twice-a-day potentiates markers of mitochondrial biogenesis in men"

**Supplementary Table 1 | Details of PCR primers used for RT-qPCR**

| Gene | Primer Efficience (%) | Forward Sequence | Reverse Sequence |
| --- | --- | --- | --- |
| PGC-1α | 103.6 | 5´-CAGCCTCTTTGCCCAGATCTT-3´ | 5´-TCACTGCACCACTTGAGTCCAC-3´ |
| PGC-1α1 | 102.9 | 5´-ATGGAGTGACATCGAGTGTGCT-3´ | 5´-GAGTCCACCCAGAAAGCTGT-3´ |
| PGC-1α4 | 111.9 | 5´-TCACACCAAACCCACAGAGA-3´ | 5´-TCACACCAAACCCACAGAGA-3´ |
| COX IV | 103.6 | 5´-GAGCAATTTCCACCTCTGC-3´ | 5´-CAGGAGGCCTTCTCCTTCTC-3´ |
| CPT1 | 111.0 | 5´-ACAGTCGGTGAGGCCTCTTA-3´ | 5´-CCACCAGTCGCTCACGTAAT-3´ |
| NDUF | 109.0 | 5´-TCAGATTGCTGTCAGACATGG-3´ | 5´-TGGTGTCCCTTCTATCTTCCA-3´ |
| SDH | 105.0 | 5´-AAATGTGGCCCCATGGTATTG-3´ | 5´-AGAGCCACAGATGCCTTCTCTG-3´ |
| Cytochrome C | 98.8 | 5´-GGGCCAAATCTCCATGGTCT-3´ | 5´-TCTCCCCAGATGATGCCTTT-3´ |
| GLUT4 | 103.6 | 5´-CTTCATCATTGGCATGGGTTT-3´ | 5´-AGGACCGCAAATAGAAGGAAGA-3´ |
| β-HAD | 80.6 | 5´-TGGACAAGTTTGCTGCTGAACAT-3´ | 5´-TTTCATGACAGGCACTGGGT-3´ |
| CD-36 | 119.0 | 5´-TTGATTGAAAAATCCTTCTTAGCCA-3´ | 5´-TGGTTTCTACAAGCTCTGGTTCTT-3´ |
| PFK | 97.6 | 5´-AAGACATCAAGAATCTGGTGGTTA-3´ | 5´-TCCAAAAGTGCCATCACTGC-3´ |
| CS | 113.5 | 5´-TGGGGTGCTGCTCCAGTATT-3´ | 5´-CCAGTACACCCAATGCTCGT-3´ |
| PPARα | 92.7 | 5´-GGCAGAAGAGCCGTCTCTACTTA-3´ | 5´-TTTGCATGGTTCTGGGTACTGA-3´ |
| PPARɣ | 109.0 | 5´-CTTGTGAAGGATGCAAGGGTT -3´ | 5´-GAGACATCCCCACTGCAAGG -3´ |
| UCP3 | 89.5 | 5´-CCACAGCCTTCTACAAGGGATTTA-3´ | 5´-ACGAACATCACCACGTTCCA-3´ |
| Tfam | 109.3 | 5´-CCGAGGTGGTTTTCATCTGT-3´ | 5´-GCATCTGGGTTCTGAGCTTT-3´ |
| PDK4 | 99.7 | 5´-GCAGCTACTGGACTTTGGTT-3´ | 5´-GCGAGTCTCACAGGCAATTC-3´ |
| p53 | 101.8 | 5´-GTTCCGAGAGCTGAATGAGG-3´ | 5´-TTATGGCGGGAGGTAGACTG-3´ |
| PPARβ/δ | 103.7 | 5´-CATCATTCTGTGTGGAGACCG-3´ | 5´-AGAGGTACTGGGCATCAGGG-3´ |
| TFEB | 102.0 | 5´-CAGATGCCCAACACGCTACC-3´ | 5´-GCATCTGTGAGCTCTCGCTT-3´ |
| CHCHD4 | 107.0 | 5´-GCTTGGCTGTTCCTTGTTATTC -3´ | 5´-GTTTCCTCTCTTGCTGCTACTC -3´ |
| p21 | 99.8 | 5´-GCAGACCAGCATGACAGATTT -3´ | 5´-GATGTAGAGCGGGCCTTTGA -3´ |
| GAPDH | 106.0 | 5´-AATCCCATCACCATCTTCCA-3´ | 5´-TGGACTCCACGACGTACTCA-3´ |
| B2M | 98.0 | 5´-TGCTGTCTCCATGTTTGATGTATCT-3´ | 5´-TCTCTGCTCCCCACCTCTAAGT-3´ |
| TBP | 99.0 | 5´-CAGTGACCCAGCAGCATCACT-3´ | 5´-AGGCCAAGCCCTGAGCGTAA-3´ |
| Cyclophilin | 100.0 | 5´-GTCAACCCCACCGTGTTCTTC-3´ | 5´-TTTCTGCTGTCTTTGGGACCTTG-3´ |
| 18S | 99.0 | 5´-CTTAGAGGGACAAGTGGCG-3´ | 5´-GGACATCTAAGGGCATCACA-3´ |
| ACTB | 107.0 | 5´-GAGCACAGAGCCTCGCCTTT-3´ | 5´-TCATCATCCATGGTGAGCTGGC-3´ |

PGC- 1α, peroxisome proliferator-activated receptor-ɣ coactivator 1α; COX IV, cytochrome c oxidase subunit IV; CPT1, carnitine palmitoyltransferase 1; NDUF, NADH:ubiquinone oxidoreductase; SDH, succinate dehydrogenase; GLUT4, Glucose transporter type 4; β-HAD, 3-hydroxyacyl-CoA dehydrogenase; CD36, fatty acid translocase cluster of differentiation 36; PFK, phosphofructokinase; CS, citrate synthase; PPARα, peroxisome proliferator-activated receptor alpha; PPARɣ, peroxisome proliferator-activated receptor delta; UCP3, uncoupling protein 3; Tfam, mitochondrial transcription factor A; PDK4, pyruvate dehydrogenase kinase 4; p53, p53 protein; PPARβ/δ, peroxisome proliferator-activated receptor beta/delta; TFEB, transcription elongation factor; CHCHD4, coiled-coil-helix-coiled-coil-helix domain containing 4; p21, p21 protein; GAPDH, glyceraldehyde 3-phosphate dehydrogenase; B2M, β-2-microglobulin; TBP, TATA-box binding protein; 18S, 18S ribosomal RNA; ACTB, actin beta.
